# Role of Anthropogenic drivers in altering the forest community structure in a prime tiger habitat in central India

**DOI:** 10.1101/2021.04.08.439106

**Authors:** Soumya Dasgupta, Tapajit Bhattacharya, Prafulla Bhamburkar, Rahul Kaul

## Abstract

Tropical forests are complex systems with heterogenous community assemblages often threatened under conservation conflicts. Herbivory and disturbances affect the diversity and species assemblages within forest patches having different disturbance regimes. We studied the change in plant community composition and structure under a disturbance gradient in the tropical dry deciduous forest of the corridor area between Nagzira-Navegaon Tiger reserve of central India. We tested the hypothesis that the plant community will change along the proximity gradient from the human settlement depending on the anthropogenic stress. We sampled 183 nested quadrat plots to collect data on species abundance and various disturbance parameters. Density, diversity, and Importance Value Index were calculated from the collected data on species abundance and girth at breast height (GBH) of individual tree species. We did multivariate analysis to assess the changes in species assemblage along the disturbance gradients. We found 76% dissimilarity between the plant communities in the three disturbance gradients from near to far from the villages perpetrated by the difference in mean abundance of species like *Tectona grandis, Terminalia sp*, and *Largerstroemia parviflora*. The anthropogenic factors significantly influence the density and diversity of tree species and regeneration classes. We found the abundance of regeneration class increased along the distance from the villages. The study intensifies the need for proper management and conservative approach to preserve the minimum diversity of the forest patches for its structural and functional contiguity as a corridor in the central India’s highly susceptible and intricate corridor framework.

## Introduction

The earth is undergoing rapid environmental changes because of human actions (Pimm et al. 1995; Vitousek et al. 1997; Tilman et al. 2001). From the ages of human civilization, settlement has been the primary human footprint on the terrestrial ecosystem. Human settlement in a natural or seminatural habitat disrupts the natural system, and exposes a wildland-urban interface (Bar-Massada et al. 2014). Within the interface, there exist two-way interactions between the human and natural ecosystem, and the extent of these interactions occurs deep in the surrounding natural landscape (Radeloff et al. 2005a). In the wildland-urban interface, there are no demarcated boundaries between infrastructure and wildland, causing conversion of natural habitats into other landcover types as well as fragmentation of contiguous habitat into smaller fragments (Bar-Massada et al. 2014; Fahrig, 2003). Through introduced barriers of different landcover types, fragmentation disrupts the ability of plant and wildlife populations to disperse and regenerate after physiological and demographic disturbance (Hanski and Ovaskainen 2000; Edwards et al. 2004). It also increases the number of open edges in the habitat further and increases the probability of disturbance in the remnant vegetation patches (Wiens 1992).

Forests are an enormously complex system with highly heterogeneous plant communities that are important to harbor high biodiversity and maintain ecosystem balance (Richards 1996; Majila and Kala 2010; Kala 2015). Forest attributes like tree composition, community structure, and diversity patterns are affected by environmental as well as anthropogenic variables acting on it over a considerable time. (Armesto and Picket 1985; Tilman et al. 1997; Karanth et al. 2006; Timilsina et al. 2007; Gairola et al. 2008; Ahmed et al. 2010; Somanathan and Borges 2000; Bagchi and Ritchie 2010; Bagchi et al. 2012). Although forest structure and composition change over time as a natural successional process (Clement and Junquiera, 2010), ever-increasing anthropogenic pressure has accelerated the rate of change (Kala and Dubey 2012). Fragmented and altered forest structures with disrupted ecosystem processes are more prone to climate hazards (Noss 2001), thus, they are in focus for immediate conservation initiatives by the resource managers for maintaining ecological resilience. (Kala and Dubey 2012).

Recognized for the richest biological communities extant on the earth, tropical forests harbor a significant proportion of global biodiversity (Myers et al. 2000; Baraloto et al. 2013). Tropical dry forests constitute 17% of currently standing tropical forests (UNEP-WCMC Forest Programme 2011), and support large populations of forest-dependent people (Miles et al. 2006). Tropical forests provide ecosystem services such as species conservation, prevention of soil erosion, carbon sequestration, and habitat preservation for plants and animals (Armenteras et al. 2009). The long history of human settlement and exposure to the natural forest has already influenced most of the forest habitat’s structure and community composition considered to be ‘natural’ (Heywood and Iriondo 2003). Anthropogenic disturbances like overgrazing of livestock, logging, encroachment, and conversion of forest land for agriculture and settlement, are the major causes for habitat conversion (Kothari *et al*. 1989; van Schaik *et al*. 1997; Sekercioglu, 2002; Sinclair *et al*., 2002; Saberwal & Rangarajan 2003; Millennium Ecosystem Assessment, 2005; Chown, 2010; Anitha et al. 2010). Extraction of forest biomass in the form of fodder and fuelwood collection, extraction of timber and non-timber forest product (NTFP), and grazing are the most widespread pressure in the tropical forest, as the dependency of rural populations in developing countries for their livelihood and sustainability is very high (Pattanayak *et al*. 2003; Agarwala 2016). As more land for agriculture and more pressure on timber and NTFP is expected with an increasing population, more pressure on forest is intuitive (Arnoldo 2000). At the current rate of deforestation and degradation of natural habitat, about 20 and 50 percent of tropical forests are expected to be lost (Wilson 1989).

Tropical moist deciduous, dry deciduous, and evergreen forests account for 72 percent of the 64 Mha forest cover area of India (Ravindranath and Sukumar 1998). The last few decades have witnessed an increasing rate of deforestation of tropical forests throughout the world and its associated biodiversity has also been depleted (Ravikanth et al 2000). India has the largest livestock population globally, and the number increased from 429.9 to 500 million from 1987 to 2000. It is expected to exceed 5000 million by the end of 2010 and of these 270 million grazes on forest land, leading to degradation of forest structure and regeneration (Anon 2002; Ali et al 2009). Anthropogenic disturbances like grazing and tree harvesting influence the regeneration of woody species and affect the vegetation composition of the forest (Colter 2006). The change in vegetation structure and tree composition significantly impacts the native faunal populations, with specialized niche requirements (Bawa & Seidler 1998).

The tropical dry deciduous forest of central India supports a large number of protected areas interspersed within areas having dense human population, which acts as a barrier for the dispersal of major faunal species like the Tiger and its prey species. Forest patches outside the protected areas experience high anthropogenic pressure. The present study is a part of the Central India Tiger Habitat Conservation project initiated by Wildlife Trust of India in the corridor area between the Nagzira-Navegaon Tiger Reserve of the Eastern Vidarbha region of Maharashtra. There are more than 100 villages within the corridor area, and the villagers are dependent on the forest for its ecosystem services. The impact of human disturbances is also high, as the forest area is used for Tendu (*Dyospyros melanoxylon*) and Mahua (*Madhuca longifolia*) collection from the month of March to June (Lal 2012, Chavan et al. 2016). We aimed to detect the community composition and identify factors affecting the changes in the plant community in different anthropogenic disturbance gradients. To estimate the impact of anthropogenic influence, the hypothesis assumed that proximity of the forest site to human settlement is more disturbed than the distant sites depending on the fact that harvesting cost increases with distance from the human settlement (Uma Shaankar et al. 2003). We also assessed the factors associated with the regeneration of different species and the difference in regeneration patterns along the disturbance gradient. The study is crucial to establish a baseline and assess the dynamics of vegetation pattern in the dry deciduous forest patches of the Central Indian Landscape.

### Study area

We carried out the study in the Gondia forest division of Maharashtra (Figure 1). The division is situated between 20°39’ and 21°38’ North latitude and 79°50’ and 80°41’ East in between the Central Indian Tiger Landscape. The Tiger reserves and the forest divisions in the Central Indian landscape are globally important tiger habitat conservation landscapes (Wikramanayake et al 1998). The National Tiger Conservation Authority (NTCA) also recognizes the importance of this landscape for long-term tiger conservation (Jhala et al 2008). The study area acts as a corridor between Nagzira-Navegaon Tiger reserve and Kanha Tiger Reserve and Navegaon National Park and plays a significant role as the buffer of the Nagzira Navegaon Tiger Reserve. The study area in Nagzira Navegaon Corridor has a varied management regime, as it is outside the Tiger reserve. The forest area is 1731.78 Km^2^ comes under the Gondia Forest Division, except the areas allotted to Forest Development Corporation of Maharashtra (FDCM) limited, where both conservation and economic benefits are associated. There are more than 100 villages with varied population density situated within the area. Major villages congregated near Jambdi and Shenda. National Highway 6 and state highway 37 is crossing through the area connecting Nagpur to the state of Chattisgarh and Goregaon, Sadak Arjuni, Deori Tehsil, and Bhandara district, respectively. The population residing in the corridor area depends on the forest not only for fuelwood but to a varied extent on Tendu and Mohua collection from late February to early June. The economy of more than 50% of the residents in the corridor depends largely on the collection and selling of Tendu leaves and Mohua flowers and their bi-products during that season (Lal 2012, Chavan et al. 2016). The pressure of domestic livestock and the free-ranging cattle population also exerting pressure on the forest vegetation. Fire incidents are also frequent during the end of the summer season and sometimes induced by the Mohua and Tendu collectors. The fire incidences before the Mohua flowering clear the forest floor and helps in the collection of Mohua flower from the clear forest floor and it also helps the tender leaves of Tendu to grow, which follows the Mohua collection season. The area is having hilly terrain with moderate to steep slopes dissected by rainfed streams. The climate of the area is generally hot and dry with relatively cold winter months from November to February. The average annual rainfall is about 1200mm and major rainfall is received from June to September. The main rivulets traversing the area are Kanhan, Chulband, Gavri, and Bagh. The forest type belongs to the tropical dry deciduous forest (5A) as per Champion and Seth (1968). Garari (*Cleistanthus collinus*) is the dominant species with Teak (*Tectona grandis*), Bija (*Pterocarpus marsupium*), Saja (*Terminalia tomentosa*), Surya (*Xylia xylocarpa*), and Dhaora (*Anogeissus latifolia*) being the other valuable timber species. Dongargaon and Navegaon have been the main timber depot of the division and firewood also sold from the depots at Deori, Chichgarh, Kohmara, etc. The division is also known for bamboo and Tendu production and has about 29 tendu units fetching crores of rupees as revenue. The forests face severe grazing pressure due to concentrated grazing by cattle including a large number of goats and sheep. As per the data available from the census 2011, 62 % of the population of Gondia district lives in rural areas. The cattle census data 2012 reveals the presence of 1.32 million of domestic animals in the district.

**Figure 1 –.**
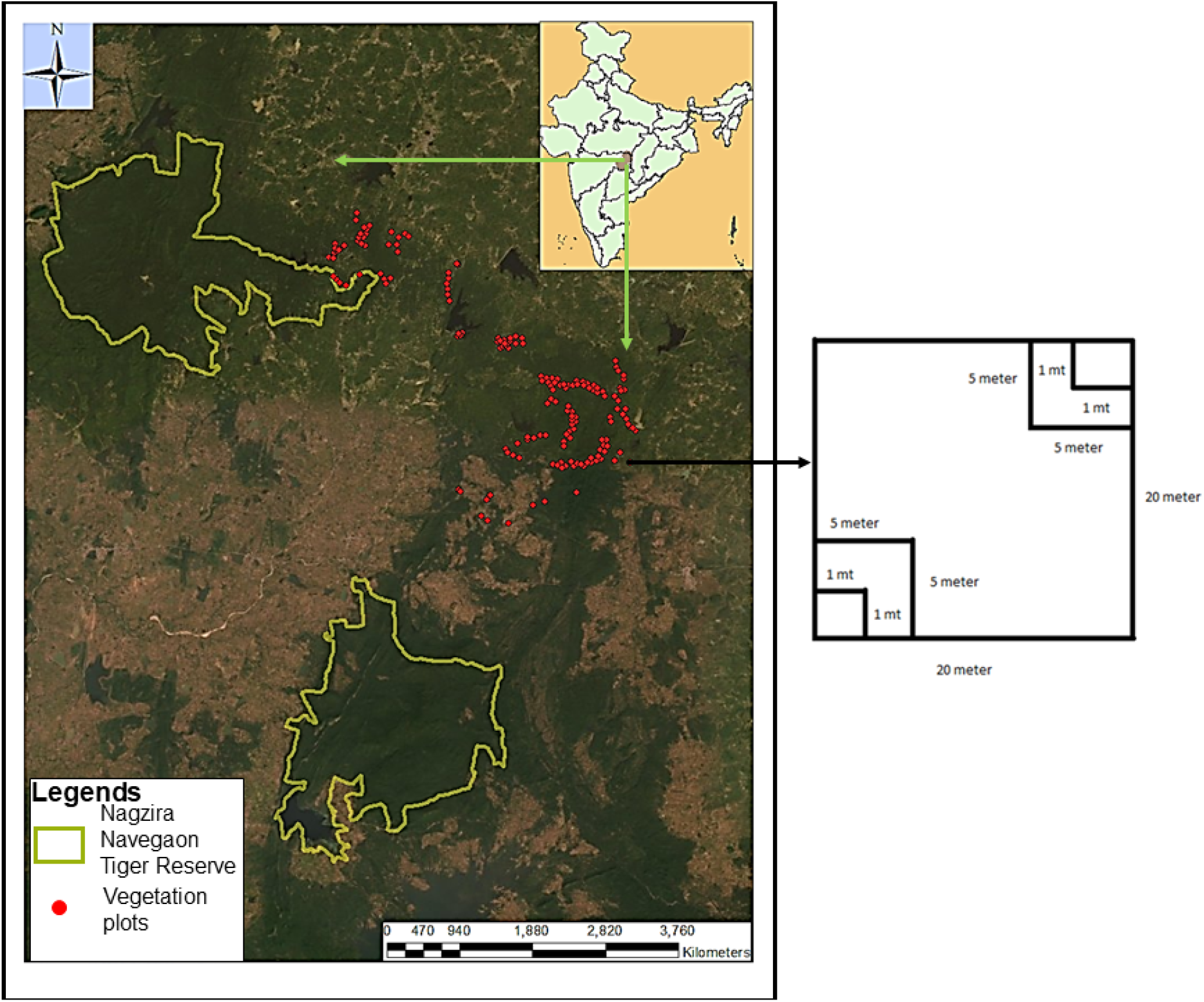
Study area in Nagzira Navegaon corridor showing the location of vegetation plots and diagrammatic scheme of nested quadrat for vegetation sampling

### Methodology

We surveyed the forested areas of Nagzira Navegaon Corridor, within Goregaon, Sadak Arjuni, and North Deori Range of the Gondia Forest division and Jambadi I and Jambadi II range of FDCM during the study. We did vegetation sampling following the nested quadrate method and maintained a minimum 200-meter distance within two quadrates. We took the largest plot size of 20mX20m for trees, and within that, we counted individuals of each tree species and identified them with the help of locals. Later, we confirmed them by consulting the taxonomists and subject specialists. Within the 400m^2^ plot, we laid two 5mX5m plots in the diagonally opposite direction to sample shrub species and regeneration classes. We counted each shrub species and saplings and recorded their basal diameter and height. Within each 25m^2^ plot, we laid one 1mX1m plot to sample herbs and seedlings (Figure 1). The tree cover, shrub cover, and herb were measured using the line intercept method. We laid a total of 183 vegetation plots within the study area. The girth at breast height (1.3 meters) and the height of each tree species were measured. For each shrub species and regeneration class, we measured the height and basal diameter. We collected data on the disturbance parameters like the presence of trail, presence of grazing sign, presence of logging and lopping, and any other anthropogenic pressure. The presence of animal signs and tracks within the vegetation plots were recorded.

We divided the vegetation plots into three clusters according to the distance from the nearest villages to asses the differences in the vegetation structure in the three-distance gradient from near to far of the villages. The distance gradients were denoted as one (0-1 km from the villages), two (1-2 km from the villages), and three (2-5 km from the villages). We estimated different species richness estimators (Chao 1 and 2, Jacknife 1, Jacknife 2, and Bootstraps), and generated species accumulation curves to assess the completeness of the sampling. The estimation of Shanon diversity index, Simpson index, evenness index, and Fisher’s alpha was done. We estimated the relative density, relative abundance, and relative dominance for different species recorded from the vegetation plots and enumerated the Importance value index (IVI, Nguyen et al. 2015) for each species. We also enumerated the average canopy height, average girth at breast height (GBH), tree density, diversity, sapling density, and diversity for plot-wise and in total from the collected data for each distance gradient. The difference among the means of the parameters in three distance gradients were also estimated using Kruskal Wallis test (Kruskal and Wallis 1952) and post hoc Dune test (Dune 1961) at 95% confidence level (alpha=0.05). We did simper analysis using ‘R’ software using ‘Vegan’ package to assess the difference in community composition in three disturbance gradients and also assessed the species contributing to the differences. Permanova (Anderson 2001) was done to determine whether the communities in different distance gradients differ significantly from each other. We did the Nonmetric Multidimensional Scaling (NMDS) using ‘Bray Curtis’ distance for the plant community. The disturbance factors and vegetation structure were plotted against the plant community to assess the association of different plant species with the different disturbance factors and vegetation structure in three disturbance regimes. We analyzed the data on the regeneration class to assess the difference in regeneration patterns in different distance gradients and factors affecting the regeneration of different species in the study area. We did Generalized additive modeling (Hastie and Tibshirani, 1990) to identify the factors influencing the regeneration using ‘R’ software (R version 4.0.3, 2020-10-10). Since most of the data sets have nonnormal distributions, we used generalized additive modeling (GAM) instead of traditional linear modeling (LM). Being a classical addendum of LM, GAM provides the additivity of non-linear predictor variables of various distributions to limit the prediction of the dependent variable. GAM also provides the option of flexible specification using smooth functions that replaces the detailed parametric relationship of the covariates. We used the F value and the significance level to determine the influence of the variable.

## Result

We sampled a total of 183 vegetation plots and encountered 3293 individual trees with an average tree density of 18±1 per 400m^2^ plot. Out of the total 183 plots, 77 plots were in distance gradient one, and 56 and 50 plots are in distance gradient two and three respectively. The overall average tree diversity, canopy height, and average GBH were 6.47±0.168 cms, 12.24±0.19 meters, and 65.28±1.380 cms respectively. The average tree density varies between the plots in the three distance gradients. We calculated the density and diversity of shrubs and regeneration class in different distance gradient (Figure 2). We found significant difference in tree density, avg GBH of trees in a plot, diversity, and density of shrub and regeneration class between the three disturbance gradient, whereas, in the case of diversity of tree and height of the canopy, the difference in three disturbance gradients was not statistically significant (Details were in Supplementary table 1). The values of the Shannon index, Simpson index, Fisher’s alpha, and evenness index were 2.905, 0.905, 11.48, and 0.281 respectively. We assessed the difference in disturbance parameters like presence of cattle and trail, lopping and logging sign in the vegetation plots along the disturbance gradient. We found that all the disturbance parameters’ presence decreases along the gradient except the presence of trail (Figure 3). The presence of trail increases in disturbance gradient two from that of one, although decreases in disturbance gradient three.

**Figure 2.**
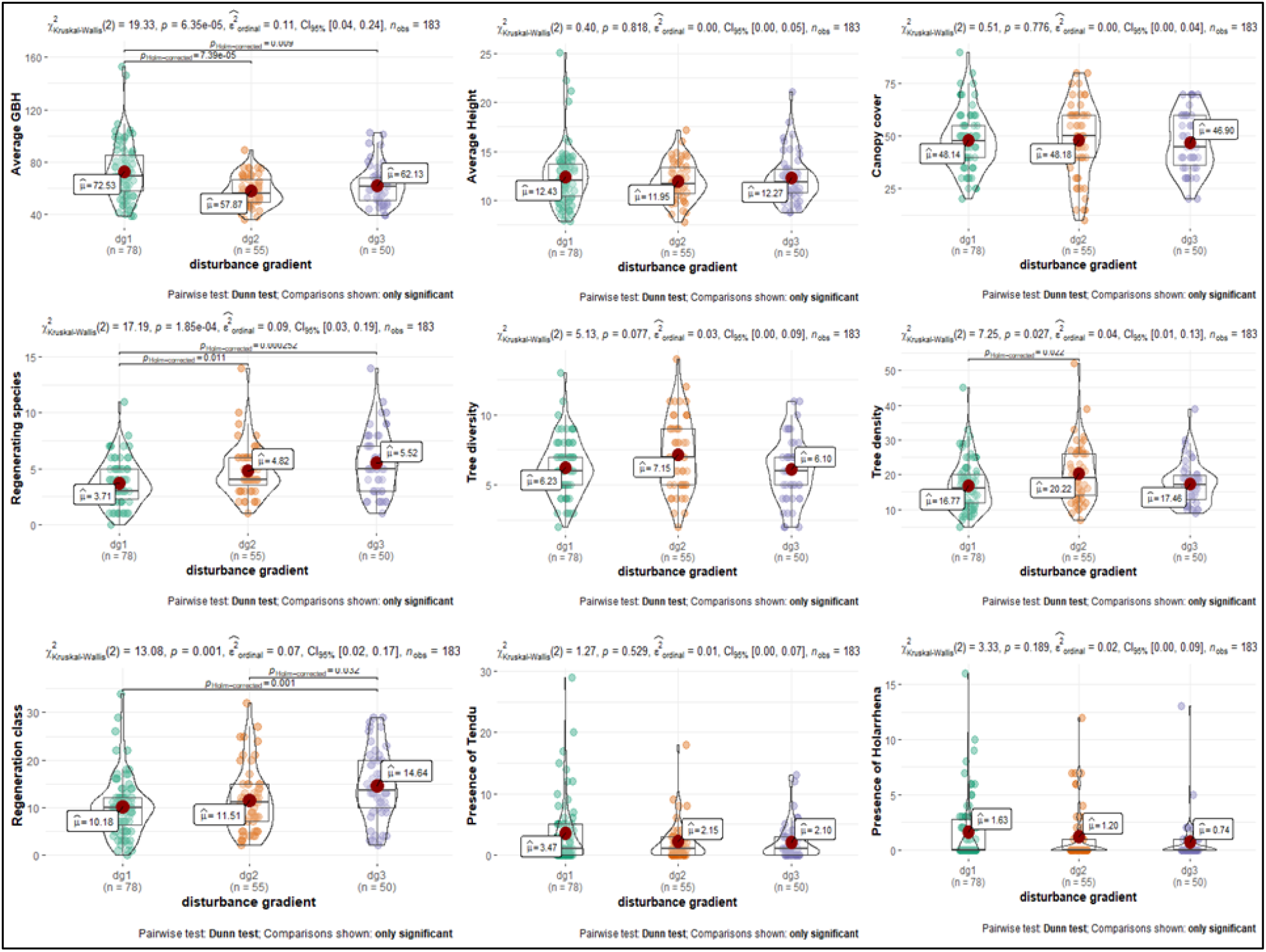
Box-violineplot showing comparative values of Average GBH, Average canopy height, Canopy cover, Tree diversity, Tree density, Regeneration class, Regenerating species diversity and presence of Tendu and presence of Holarrhena sp in three disturbance gradients from near to distantly placed from villages denoted as dg1, dg2 and dg3. The Value of Kruskal Wallis Chi square test and post hoc Dune test were given for each variables.

**Figure 3.**
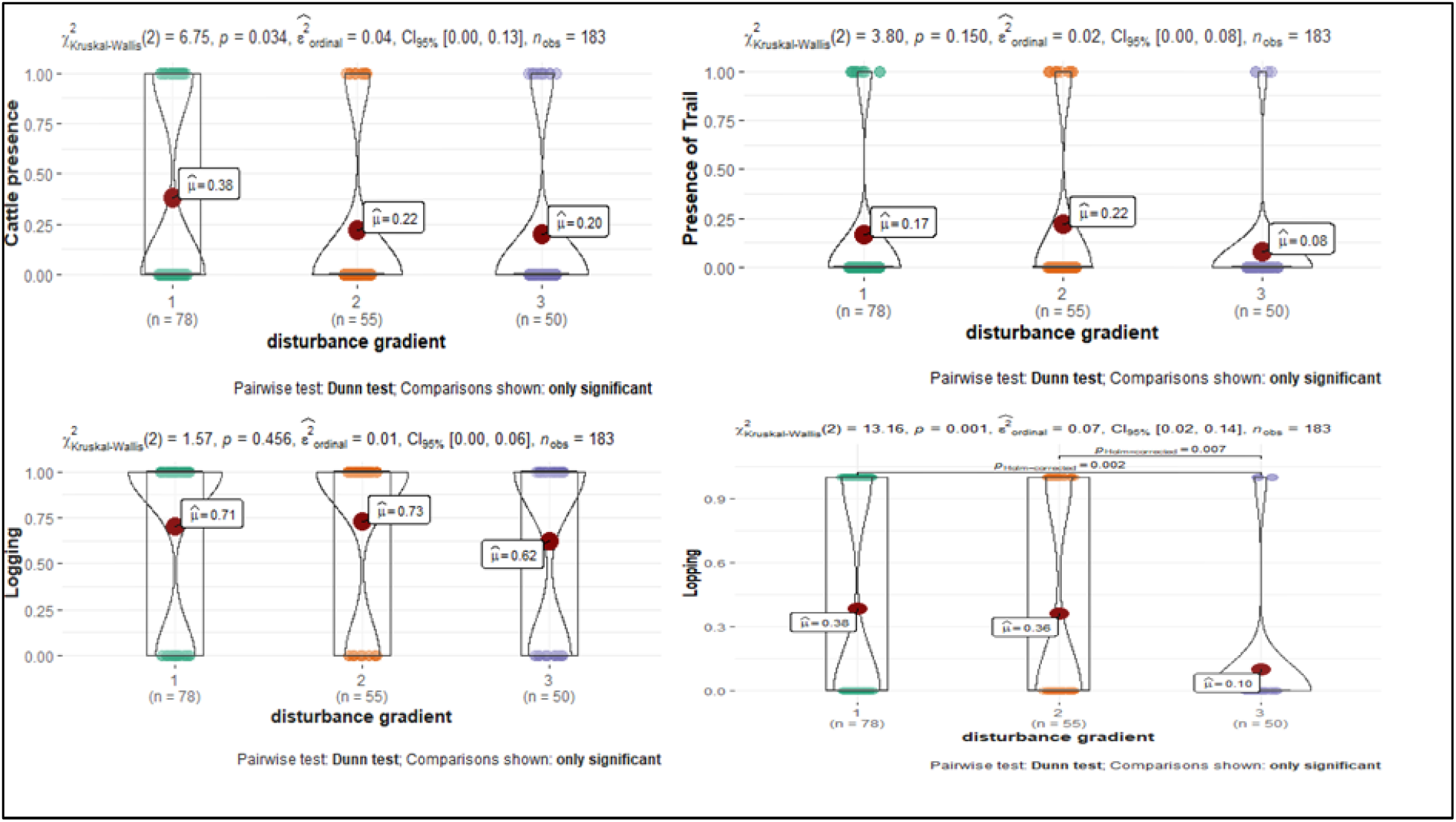
Box-violineplot showing comparative presence absences of different disturbance parameters (Cattle presence, Presence of Trail, Logging, and Lopping) in the three disturbance gradient. The Value of Kruskal Wallis Chi square test and post hoc Dune test were given for each variables.

We recorded 61 plant species during the survey of which 50 species were recorded from distance gradient two and 46 and 45 species recorded from distance gradient three and one. The species accumulation curve shows moderate to high completeness of species accumulation during the sampling (Figure 4). The results of the different diversity estimators were given in table 1.

**Figure 4 –.**
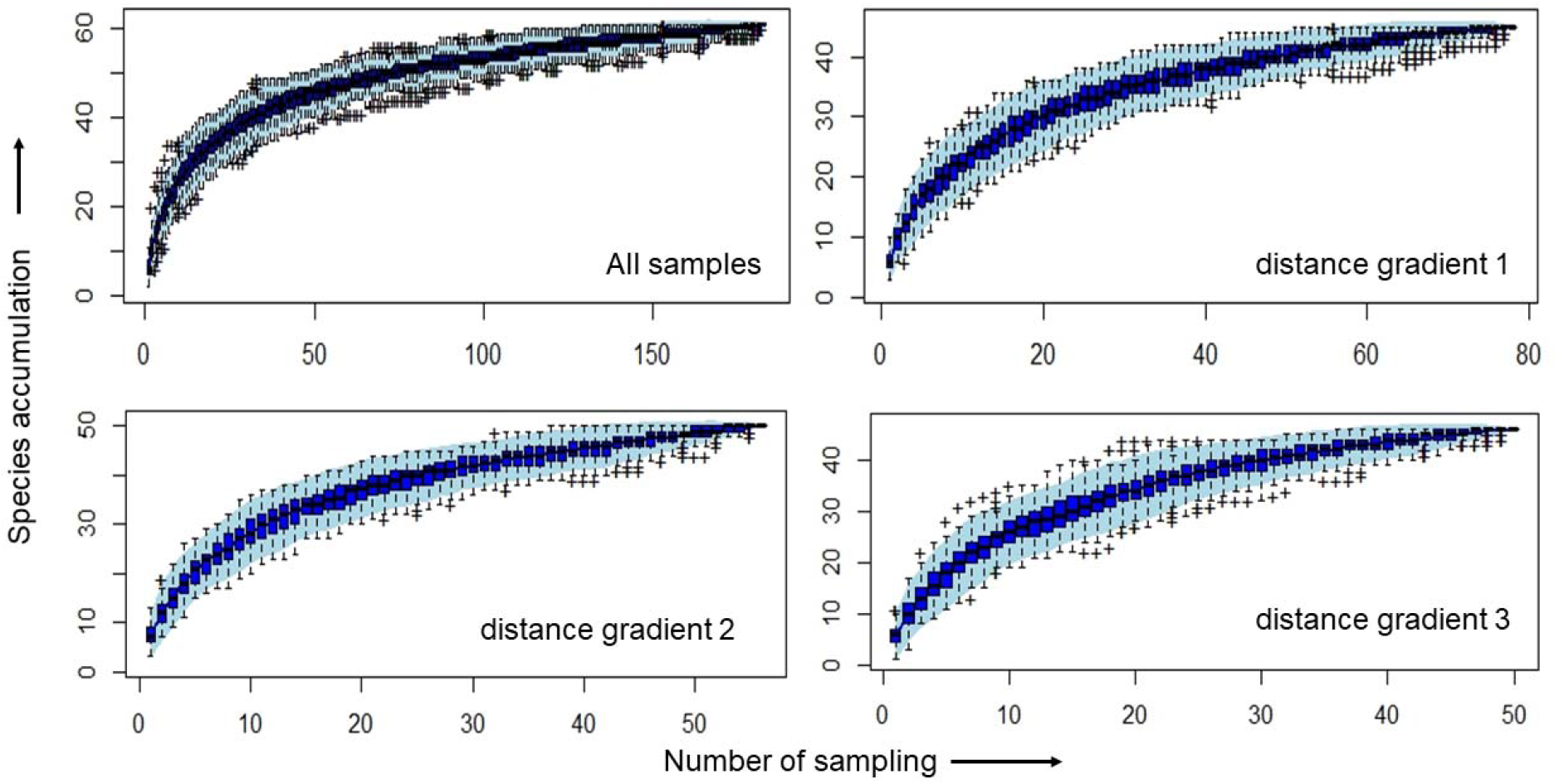
Species accumulation curve for overall sampling and sampling in three disturbance gradients on accordance to distance from nearest villages.

**Table 1 –.** Different diversity estimators in different distance gradient

|  | species | Chao | Jackknife 1 | Jackknife 2 | Bootstrap | Sampling unit |
| --- | --- | --- | --- | --- | --- | --- |
| Overall | 61 | 93.488±26.19 | 74.92±3.98 | 85.819 | 66.76±2.103 | 183 |
| Distance gradient one | 45 | 53.22±6.82 | 54.87±3.42 | 58.845 | 49.795±2.1 | 77 |
| Distance gradient two | 50 | 57.857±5.94 | 61.78±3.93 | 64.836 | 55.69±2.411 | 56 |
| Distance gradient three | 46 | 51.44±4.53 | 55.8±3.677 | 56.936 | 51.168±2.291 | 50 |

Teak (*Tectona grandis*) was the dominant species found with the highest IVI value (50.462) followed by Saja (*Terminalia tomentosa*), Sihina (*Lagerstroemia parviflora*), Bija (*Pterocarpus marsupium*), Dhaora (*Anogeissus latifolia*). Table 2 shows the top ten species with the highest IVI value.

**Table 2 –.** Contribution of different species (top ten) for overall dissimilarity between species assemblages in different plant communities in three disturbance gradients from near to distantly placed from villages along with the IVI values of the tree species.

| Tree | distance gradient one and two |  | distance gradient one and three |  | distance gradient two and three |  | Mean abundance of tree species |  |  | IVI |  |
| --- | --- | --- | --- | --- | --- | --- | --- | --- | --- | --- | --- |
| Species | Av. Dissimilarity | Contribution % | Av. Dissimilarity | Contribution % | Av. dissimilarity | Contribution. % | One | Two | Three | Value | Rank |
| *Tectona grandis* | 14.18 | 19.12 | 15.13 | 19.47 | 14.65 | 19.61 | 2.52 | 4.73 | 4.74 | 50.46259 | 1 |
| *Terminalia tomentosa* | 8.927 | 12.04 | 8.404 | 10.81 | 8.213 | 11 | 2.91 | 3.16 | 1.32 | 39.52649 | 2 |
| *Lagerstroemia parviflora* | 7.044 | 9.502 | 6.709 | 8.632 | 5.644 | 7.557 | 2.17 | 1.93 | 1.28 | 27.50358 | 3 |
| *Anogeissus latifolia* | 5.566 | 7.509 | 5.316 | 6.839 | 3.856 | 5.163 | 1.65 | 1.14 | 0.66 | 21.08546 | 5 |
| *Buchanania lanzam* | 4.425 | 5.969 | 3.318 | 4.269 | 4.005 | 5.362 | 0.948 | 1.46 | 0.622 | 16.40634 | 6 |
| *Cleistanthus collinus* | 4.298 | 5.799 | 7.458 | 9.596 | 6.26 | 8.382 | 1.01 | 0.696 | 2.04 | 15.51898 | 7 |
| *Pterocarpus marsupium* | 3.738 | 5.043 | 4.691 | 6.036 | 3.798 | 5.085 | 1.05 | 0.786 | 1.2 | 21.24336 | 4 |
| *Lannea coromandelica* | 2.823 | 3.808 | 2.614 | 3.364 | 1.805 | 2.416 | 0.766 | 0.589 | 0.32 | 12.72077 | 8 |
| *Madhuka latifolia* | 2.614 | 3.526 | 2.057 | 2.646 | 2.145 | 2.872 | 0.519 | 0.768 | 0.3 | 11.59404 | 9 |
| *Xylia xylocarpa* | 2.347 | 3.166 | 3.18 | 4.092 | 4.459 | 5.971 | 0.0779 | 0.804 | 1.04 | 7.89958 | 10 |

The simper analysis result showed an overall average dissimilarity within the three distance gradients as 75.53% and the dissimilarity between plant communities of distance gradient one and two as 74.12% and that of between distance gradient one and three as 77.72%. The difference in plant communities between distance gradient two and three was 74.69%. The tree species contributing to major dissimilarity for plant communities one and two, one and three, and two and three are given in supplementary Table 2 respectively (Supplementary table 2a for gradient 1 and 2, 2b for gradient 1 and 3 and 2c for gradient 2 and 3).

The result of simper and IVI showed that the difference in the community structure is because of the differed abundance value of the major species that came out as per IVI values. Major species which are responsible for the dissimilarity in community composition are *Teak* (*Tectona grandis*), *Saja* (*Terminalia tomentosa*), Sihina (*Lagerstroemia parviflora*), Garadi (*Glestanthus collinus*), Dhaora (*Anogiessus latifolia*), Char (*Buchanania lanzam*), Bija (*Pterocarpus marsupium*), Mohoi (*Lannea coromandelica*), Mohua (*Madhuka lattifolia*), Surya (Xylia xylocarpa). Permanova result shows a significant difference in community structure within the three disturbance sites, both for tree species and regeneration classes. (Table 3)

**Table 3 –.** Permanova results for tree species assemblages and assemblages of regeneration class among the three disturbance gradients.

| Tree species | Df | Sum of<br>Sqs | Mean<br>Sqs | F.<br>Model | R2 | Pr(>F) |  |
| --- | --- | --- | --- | --- | --- | --- | --- |
| Distance gradient | 1 | 1.999 | 1.99915 | 7.0369 | 0.03742 | 0.001 | *** |
| Residuals | 181 | 51.421 | 0.28409 |  | 0.96258 |  |  |
| Totals | 182 | 53.42 |  |  | 1 |  |  |
| Regeneration<br>Class |  |  |  |  |  |  |  |
| Distance gradient | 1 | 2.083 | 2.08324 | 6.3544 | 0.0341 | 0.001 | *** |
| Residuals | 180 | 59.012 | 0.32784 |  | 0.9659 |  |  |
| Totals | 181 | 61.095 |  |  | 1 |  |  |
| Significance<br>codes | 0 '***' | 0.001 '**' | 0.01 '*' | 0.5 '.' | 0.1 ' ' | 1 |  |

Two-dimensional Nonmetric multidimensional scaling plot on species abundance data showed the diverse species distribution in three disturbance gradients (Figure 5). Species such as Saja (*Terminalia tomentosa*), Hidda (*Terminalia chebula*), Char (*Buchanania lanzam*), Sihina (*Lagerstroemia parviflora*), Mahua (*Madhuca lattifolia*), Palas (*Butea monosperma*), Bhera (*Chloroxylon swietenia*) formed distinct assemblages. Teak (*Tectona grandis*) and Garadi (*Clestanthus collinus*) form distinct distribution separately, whereas we found Surya (*Xylia Xylocarpa*) only in the sampling plots far away from the villages.

**Figure 5 –.**
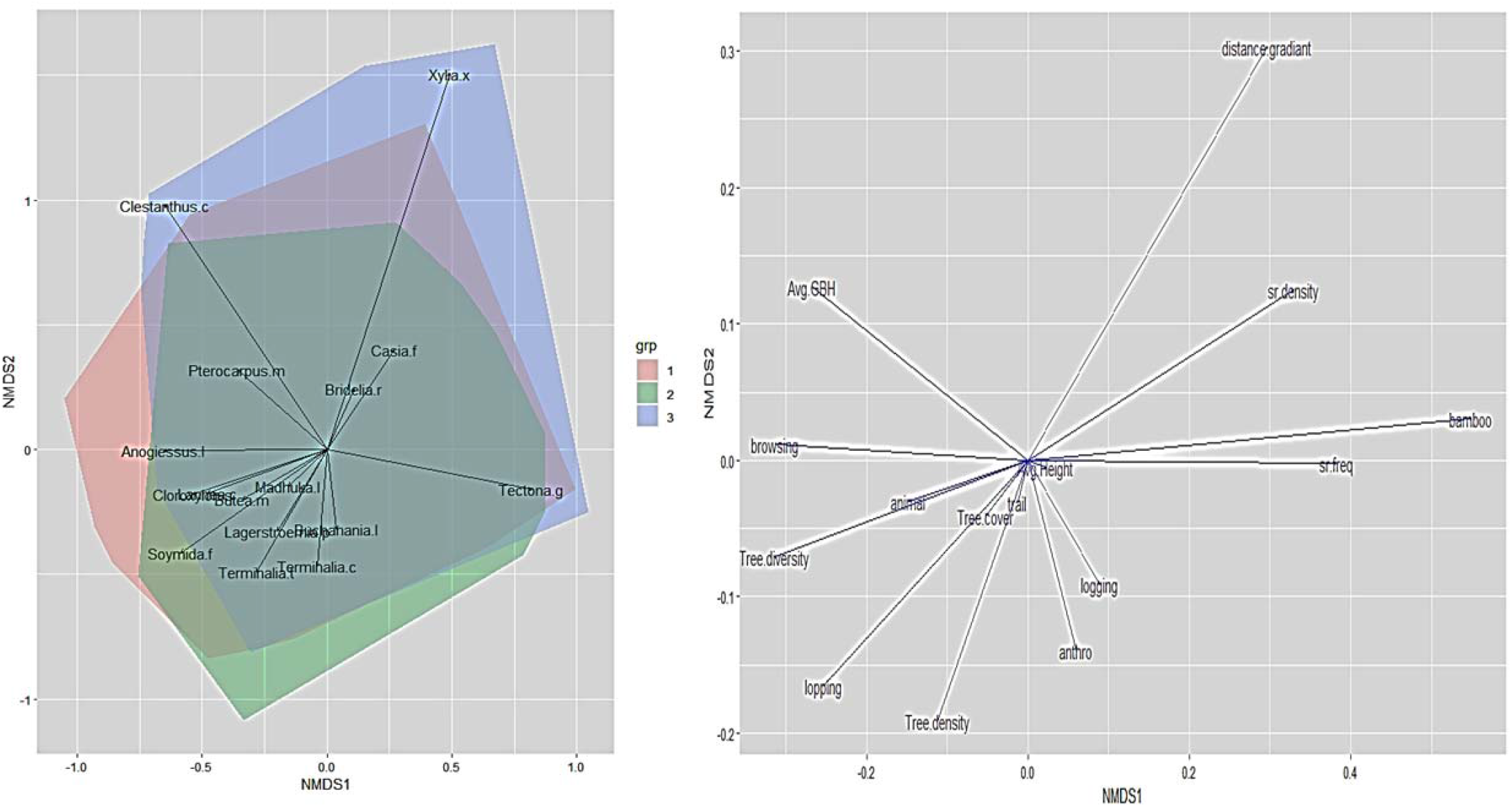
NMDS plot with values of Ward’s distance of environmental variables and species assemblages in three disturbance gradients with NMDS 1 in X axis and NMDS 2 in Y axis.

We used disturbance parameters such as lopping sign, logging sign, cattle grazing and browsing evidence, other anthropogenic pressure and parameters like tree density, tree diversity, distance gradient, average GBH of the trees, density, and diversity of regeneration class, canopy cover data as the biophysical variable for NMDS plot. Anthropogenic pressure and logging negatively affected tree diversity, average canopy height, and average GBH of the trees found within a plot. We found distance gradient is negatively attributed to the tree density and animal presence. As expected, our result shows anthropogenic pressure was high near the villages than the faraway areas.

Regeneration and recruitment of new trees are important for forest dynamics. We found an increasing number of species in the regeneration plot as the distance from the villages increases, so as with the total number of regenerating individuals (Figure 2). Tendu and *Holarrhena* were the major regeneration class found in all the three distance gradients, with a decreasing recruitment pattern along the distance to the villages. We found lower regeneration and recruitment of one of the economically important species Mohua (*Madhuca* sp) compared to other dominant tree species with high IVI values in the study area along the different disturbance gradient. Simper analysis shows 79.83% average dissimilarity within communities of regeneration classes of distance gradient one and two and that of between two and three and one and three were 83.15% and 79.99%. We found that more than 50% of these dissimilarities were due to the differences in the mean abundance of species like Tendu (*Diospyrous malanooxylon*), Kuda (*Holarrhena sp*), Teak (*Tectona grandis*), Saja (*Terminalia tomentosa*), Sihina (*Lagerstroemia parviflora*) and Atai (*Helicteres sp*) (supplementary table 2).

The result of Generalized additive modeling shows that the diversity of recruitment of the regeneration class was significantly influenced by distance to the villages, presence of bamboo as shade species, browsing, lopping, logging, tree density, average GBH of trees in the plot, tree cover and average height of the trees in the plot at the minimum 95% significance level. (table 4). The total number of recruitments in a particular plot was attributed by distance to villages and the presence of bamboo at the 95% significance level (Table 4).

**Table 4 –.** Factor influencing recruitment diversity and total recruitment of regeneration class resulted from generalized additive modeling (GAM)

| Parametric coefficients: | Recruitment diversity of regeneration class |  |  |  |  | Total recruitment of regeneration class |  |  |  |  |
| --- | --- | --- | --- | --- | --- | --- | --- | --- | --- | --- |
|  | Estimate | Std.Error | z value | Pr(>\|z\|) |  | Estimate | Std.Error | z value | Pr(>\|z\|) |  |
| (Intercept) | 2.14949 | 0.07423 | 28.959 | <2e-16 | *** | 1.238302 | 0.112219 | 11.035 | < 2e-16 | *** |
| Moderate distance | -0.06773 | 0.06609 | -1.025 | 0.305461 |  | 0.119911 | 0.099676 | 1.203 | 0.22897 |  |
| Furthest distance | 0.23455 | 0.06368 | 3.683 | 0.00023 | *** | 0.304488 | 0.097458 | 3.124 | 0.00178 | ** |
| anthropogen1 | 0.0721 | 0.06046 | 1.192 | 0.233088 |  | 0.007572 | 0.092551 | 0.082 | 0.93479 |  |
| animal1 | 0.0916 | 0.06579 | 1.392 | 0.163835 |  | -0.015054 | 0.101723 | -0.148 | 0.88235 |  |
| browsing1 | 0.20567 | 0.0731 | -2.814 | 0.0049 | ** | 0.167027 | 0.112636 | 1.483 | 0.1381 |  |
| lopping1 | -0.14372 | 0.07056 | -2.037 | 0.041671 | * | 0.008487 | 0.107691 | 0.079 | 0.93718 |  |
| trail1 | 0.11796 | 0.07074 | 1.668 | 0.095408 | . | 0.156899 | 0.103367 | 1.518 | 0.12904 |  |
| logging1 | -0.11933 | 0.06058 | -1.97 | 0.04887 | * | -0.093084 | 0.094975 | -0.98 | 0.32704 |  |
| bamboo1 | 0.21396 | 0.07205 | 2.97 | 0.002982 | ** | 0.139439 | 0.107743 | 1.294 | 0.1956 |  |
| bamboo2 | 0.51512 | 0.14109 | 3.651 | 0.000261 | *** | 0.405305 | 0.215134 | 1.884 | 0.05957 | . |
| bamboo3 | 0.79782 | 0.12607 | 6.329 | 2.47E-10 | *** | 0.6266 | 0.197138 | 3.178 | 0.00148 | ** |
| Approximate significance of smooth terms |  |  |  |  |  |  |  |  |  |  |
|  | edf | Ref.df | Chi.sq | p-value |  | edf | Ref.df | Chi.sq | p-value |  |
| s(Tree.den) | 2.915 | 3.661 | 13.736 | 0.00676 | ** | 1.813 | 2.301 | 2.471 | 0.2881 |  |
| s(Avg.GBH) | 8.624 | 8.927 | 47.372 | 1.76E-06 | *** | 6.5 | 7.588 | 12.976 | 0.0884 | . |
| s(Tree.div) | 2.308 | 2.955 | 2.931 | 0.35932 |  | 1 | 1 | 0.031 | 0.8593 |  |
| s(Avg.Ht) | 7.502 | 8.348 | 41.981 | 1.37E-06 | *** | 4.569 | 5.626 | 5.678 | 0.4078 |  |
| s(Tree.cov) | 8.506 | 8.915 | 35.923 | 3.83E-05 | *** | 1 | 1 | 1.477 | 0.2242 |  |

Signif. codes: 0 '\*\*\*' 0.001 '\*\*' 0.01 '\*' 0.05 '.' 0.1 ' ' 1

## Discussion

Most of the studies carried out on the impact of biomass extraction revealed that even a low level of resource extraction continued for a longer duration may affect the forest structure and tree composition (for example Murali *et al*. 1996; Shankar *et al*. 1998; Singh 1999; Tilman & Lehman 2001; Sagar & Singh 2004). Depending on the impact and frequencies of disturbances on the ecosystem, in some case studies, the species richness reduces (Parthasarathy 1999; Addo-Fordjou *et al*, 2009), while in some others it increases (Molino and Sabatier, 2001, Kumar and Ram 2005, Mishra *et al*., 2004, Bongers *et al*., 2009). The increase in diversity evident from some studies was in response of the intermediate level of disturbances (Intermediate disturbance hypothesis by Connell 1978) (Fox, 1979, Kumar and Ram 2005, Mishra *et al*., 2004). In our study, the diversity of tree species does not significantly differ in different disturbance gradients, although differences were significant in the species assemblages. We found that the species like Teak (*Tectona grandis*) and Garadi (*Clestanthus collinus*) form different distribution regimes. Most of the dominant species with high Importance values (IVI) like Teak (*Tectona grandis}*, Saja (*Terminalia tomentosa*), Sihina (*Lagerstroemia parviflora*), Garadi (*Glestanthus collinus*), Dhaora (*Anogiessus latifolia*), Char (*Buchanania lanzam*), Bija (*Pterocarpus marsupium*), Mohoi (*Lannea coromandelica*), Mohua (*Madhuka lattifolia*), and Surya (*Xylia xylocarpa*) have different relative abundance in the three disturbance gradient.

Anthropogenic influence on forests alters forest composition in two possible ways. It either creates conditions that increase the abundance of some species or species with high dependency for useful products to remove or replace other species (Crook and Clapp 1998; Agarwala et al. 2016). In the present study the distribution of timber species like Teak (*Tectona grandis*), Bija (*Pterocarpus marsupium*), Saja (*Terminalia tomentosa*), Surya (Xylia xylocarpa), and Dhaora (*Anogiessus latifolia*) differ in three distance gradients. Regeneration of species with high economic values, like Tendu, was high near the villages than in the distant location. The regeneration of different species and diversity of regeneration class was less near the villages. The less diversity and regeneration in the forest will also affect the future structure of the forest. We found that lopping, logging, and other anthropogenic pressure effectively shape the forest structure in the study area as the factors influence both the tree recruitment and the tree density and average GBH of the trees in the sample plots. The effect was found in the furthest distance gradient also, though it was decreasing, the persisting disturbance can affect the distribution of different plant species in the future. These poorly addressed conflicts present in the buffer of most of the protected areas present increasingly difficult obstacles for the effective conservation and management of many conservation priority areas. Conservation conflicts often serve as proxies for the conflicts over more fundamental, and social needs, and recognition of the values of the natural resources more often remains unrecognized.

Spatial variation in species composition is one of the most fundamental and conspicuous features of the natural world. Within a single larger community, it creates nestedness of generalist and specialist species. Herbivory and disturbances are the major factors driving the dissimilarities in species composition within a forest community. The long-term monitoring of the vegetation dynamics is necessary for the area due to persisting disturbance and the dominance of some species like Teak and Garadi was increasing. Teak was also planted by the forest department and Forest development corporation of Maharashtra (FDCM) at regular intervals for its economic benefit. On the other hand, being fire resistant Garadi has a better survival prospect than other species that are intolerant. In some of the sampling plots, Garadi was the predominant or only species found. The frequent fire incident will also support the Garadi infestation more and more in the study area. The cumulative impact will be worse and will affect the species composition as well as the community composition of this important landscape and will change the forest dynamics in the future.

Protected areas are the foundation to conserve the remaining wildlife and its habitat in the recent decades. Most protected areas (PAs) in India, are small and surrounded by ecologically unfavorable land-use regimes (Jhala et al 2008), thus unable to sustain a viable population of species with a large home range (Hanski 1994, Wikramanayake et al 1998, 2011). The establishment of secure habitat space in between the small, isolated pockets of protected areas is needed for long-term conservation of species (Weber and Rabinowitz 1996). Central Indian Landscape is a critical tiger conservation landscape (Sanderson et al 2006), with more than 10 Tiger reserves interconnected with thin linkages of corridors. The corridors connecting the Tiger reserves are essential for structural and functional connectivity. The study area falls in the corridor of Nagzira Navegaon Tiger reserve and the Kanha Navegaon Tiger corridor through the Balaghat Hill forests of Madhya Pradesh. So, the structural integrity of the forest is important for the dispersal of the Tiger and other predator species like leopard and dhole populations within and between the landscape (Joshi 2013). The forest habitat structure influences the prey-predator interaction within a metacommunity. Proper management of the resources and a conservative approach is needed to preserve the minimum diversity of the forest patches for its structural and functional contiguity as a corridor in the highly sensitive and intricate corridor framework of central India.

## Conclusion

Although the study area is under different protection and conservation measures of the Gondia Forest Division and Forest Development Corporation Maharashtra (FDCM), we documented anthropogenic activity at the vegetation plots far away (2-5 km) from the villages. The difference in the tree community, tree diversity, average GBH of the trees, density, and diversity of regeneration class in different distance gradient was well attributed by the anthropogenic pressure on the forest. As the study area is within a corridor of conservation priority, both the structural and functional integrity of the forest is important. Several studies have focused on the functionality of the corridors for large carnivores in Central India and other parts of the world but along with that, the structural integrity of the forest of the corridor area also needs to be assessed, as maintenance of diversity and regeneration of the forest is equally important for the faunal population supported by the corridor forests.

## Acknowledgments

The study is a part of the Central India Tiger Habitat Conservation project of Wildlife Trust of India. The authors want to acknowledge Mr. Vivek Menon, Executive Director and CEO of Wildlife Trust of India for his support and encouragement throughout the study. The authors want to thank Mr. Avi Gupta, Ms. Swaroop Patankar, Ms. Tanushree Dasgupta, Mr. Gnaneshwar Raut, Mr. Hiwraj Raut, and Mr. Parag Rahangdale for assisting in data collection and data entry. The authors would like to acknowledge the Maharashtra Forest Department and Forest Development Corporation of Maharashtra (FDCM) for the permission of the study and the help during the fieldwork.

## Declaration

### Ethics approval and consent to participate

Not applicable

### Consent for publication

Not Applicable

### Availability of data and Materials

The datasets generated during and/or analysed during the current study are available from the corresponding author on a reasonable request.

## Author’s Contribution

All authors equaly contributed in study design. Data collection was done by Soumya Dasgupta. Soumya Dasgupta and Tapajit Bhattacharya analyzed the data and drafted the manuscript. All authors read and approved the manuscript.

