## Supplementary File for "Role of Anthropogenic drivers in altering the forest community structure in a prime tiger habitat in central India"

S Table 1 – Tree density, tree diversity, canopy height, Average GBH, density and diversity of regeneration class in three distance gradients.

|  | Distance gradiant 1 | Distance gradiant 2 | Distance gradiant 3 | Overall |
| --- | --- | --- | --- | --- |
| Tree density | 16.80±0.841 | 20.30±0.984 | 17.24±1.029 | 17.994±0.54 |
| Tree diversity | 6.23±0.23 | 7.145±0.34 | 6.1±0.31 | 6.47±0.168 |
| Canopy height | 12.43±0.34 | 11.95±0.26 | 12.27±0.35 | 12.24±0.19 |
| Average GBH | 72.526±2.512 | 57.867±1.534 | 62.13±2.134 | 65.28±1.380 |
| Shrub+regeneration class density | 10.307±0.69 | 11.52±0.9 | 14.58±1.032 |  |
| Shrub+regeneration class diversity | 3.82±0.21 | 4.78±0.3 | 5.76±0.433 |  |

S Table 2A - Average dissimilarity contributed by different species in distance gradient 1 and 2

| Taxon | Av. dissim | Contrib. % | Cumulative % | Mean abund. 1 | Mean abund. 2 |
| --- | --- | --- | --- | --- | --- |
| tendu | 18.19 | 22.78 | 22.78 | 3.46 | 2.22 |
| kurwa | 12.13 | 15.19 | 37.97 | 1.67 | 1.22 |
| saja | 7.518 | 9.417 | 47.39 | 0.974 | 0.964 |
| teak | 6.995 | 8.762 | 56.15 | 0.615 | 1.07 |
| sihina | 6.176 | 7.736 | 63.88 | 0.615 | 0.691 |
| atai | 5.809 | 7.276 | 71.16 | 0.128 | 1.29 |
| palsa | 5.067 | 6.347 | 77.51 | 0.615 | 0.6 |
| char | 3.836 | 4.805 | 82.31 | 0.423 | 0.491 |
| surya | 2.826 | 3.54 | 85.85 | 0.0128 | 0.8 |
| garpor | 2.257 | 2.827 | 88.68 | 0.154 | 0.218 |
| baba | 1.746 | 2.187 | 90.87 | 0.141 | 0.218 |
| behera | 1.721 | 2.156 | 93.02 | 0.282 | 0.0727 |
| tiwas | 1.575 | 1.973 | 95 | 0.0128 | 0.455 |
| mahua | 1.421 | 1.78 | 96.77 | 0.0769 | 0.218 |
| garadi | 1.038 | 1.3 | 98.07 | 0.154 | 0.0545 |
| amla | 0.5723 | 0.7169 | 98.79 | 0.0513 | 0.0364 |
| bija | 0.4202 | 0.5263 | 99.32 | 0.0256 | 0.0545 |
| aran | 0.3215 | 0.4026 | 99.72 | 0.0128 | 0.0545 |
| ghoti | 0.2231 | 0.2794 | 100 | 0.0128 | 0.0364 |

S Table 2B - Average dissimilarity contributed by different species in distance gradient 1 and 3

| Taxon | Av. dissim | Contrib. % | Cumulative % | Mean abund. 1 | Mean abund. 2 |
| --- | --- | --- | --- | --- | --- |
| tendu | 16.58 | 19.95 | 19.95 | 3.46 | 2.16 |
| kurwa | 9.809 | 11.8 | 31.74 | 1.67 | 0.76 |
| atai | 8.556 | 10.29 | 42.03 | 0.128 | 1.78 |
| teak | 8.218 | 9.883 | 51.91 | 0.615 | 1.54 |
| saja | 5.617 | 6.756 | 58.67 | 0.974 | 0.46 |
| tiwas | 5.586 | 6.718 | 65.39 | 0.0128 | 1.34 |
| surya | 5.129 | 6.169 | 71.56 | 0.0128 | 1.24 |
| palsa | 4.737 | 5.696 | 77.25 | 0.615 | 0.68 |
| sihina | 4.215 | 5.069 | 82.32 | 0.615 | 0.36 |
| char | 3.474 | 4.178 | 86.5 | 0.423 | 0.46 |
| garadi | 1.658 | 1.994 | 88.49 | 0.154 | 0.24 |
| baba | 1.52 | 1.828 | 90.32 | 0.141 | 0.24 |
| garpor | 1.481 | 1.782 | 92.1 | 0.154 | 0.12 |
| behera | 1.48 | 1.78 | 93.88 | 0.282 | 0.06 |
| ghoti | 1.381 | 1.661 | 95.55 | 0.0128 | 0.34 |
| bija | 1.262 | 1.518 | 97.06 | 0.0256 | 0.3 |
| amla | 1.154 | 1.388 | 98.45 | 0.0513 | 0.22 |
| mahua | 0.7638 | 0.9186 | 99.37 | 0.0769 | 0.08 |
| aran | 0.5235 | 0.6295 | 100 | 0.0128 | 0.1 |

S Table 2C - Average dissimilarity contributed by different species in distance gradient 2 and 3

| Taxon | Av. dissim | Contrib. % | Cumulative % | Mean abund. 1 | Mean abund. 2 |
| --- | --- | --- | --- | --- | --- |
| tendu | 12.39 | 15.49 | 15.49 | 2.22 | 2.16 |
| atai | 10.07 | 12.6 | 28.08 | 1.29 | 1.78 |
| teak | 8.859 | 11.08 | 39.16 | 1.07 | 1.54 |
| kurwa | 8 | 10 | 49.16 | 1.22 | 0.76 |
| surya | 6.639 | 8.3 | 57.46 | 0.8 | 1.24 |
| tiwas | 6.268 | 7.837 | 65.3 | 0.455 | 1.34 |
| saja | 5.294 | 6.619 | 71.92 | 0.964 | 0.46 |
| palsa | 4.377 | 5.472 | 77.39 | 0.6 | 0.68 |
| sihina | 4.128 | 5.161 | 82.55 | 0.691 | 0.36 |
| char | 3.477 | 4.347 | 86.9 | 0.491 | 0.46 |
| baba | 1.718 | 2.148 | 89.04 | 0.218 | 0.24 |
| garpor | 1.614 | 2.018 | 91.06 | 0.218 | 0.12 |
| ghoti | 1.361 | 1.702 | 92.76 | 0.0364 | 0.34 |
| mahua | 1.287 | 1.61 | 94.37 | 0.218 | 0.08 |
| bija | 1.276 | 1.595 | 95.97 | 0.0545 | 0.3 |
| garadi | 1.118 | 1.397 | 97.36 | 0.0545 | 0.24 |
| amla | 0.9657 | 1.207 | 98.57 | 0.0364 | 0.22 |
| aran | 0.6463 | 0.808 | 99.38 | 0.0545 | 0.1 |
| behera | 0.496 | 0.6202 | 100 | 0.0727 | 0.06 |
